# Neuroendocrine Prostate Cancer Drivers SOX2 and BRN2 Confer Differential Responses to Imipridones ONC201, ONC206, and ONC212 in Prostate Cancer Cell Lines

**DOI:** 10.1101/2024.08.28.610184

**Authors:** Connor Purcell, Praveen R. Srinivasan, Maximilian Pinho-Schwermann, William J. MacDonald, Elizabeth Ding, Wafik S. El-Deiry

**Affiliations:** Laboratory of Translational Oncology and Experimental Cancer Therapeutics, The Warren Alpert Medical School, Brown University, Providence, RI, 02903, USA; The Joint Program in Cancer Biology, Brown University and the Lifespan Health System, Providence, RI 02903, USA; Department of Pathology and Laboratory Medicine, The Warren Alpert Medical School, Brown University, Providence, RI, 02903, USA; Hematology-Oncology Division, Department of Medicine, Rhode Island Hospital and Brown University, Providence, RI 02903, USA; Legorreta Cancer Center at Brown University, The Warren Alpert Medical School, Brown University, Providence, RI, 02903, USA

**Keywords:** Prostate Cancer, Neuroendocrine Differentiation, ONC201, BRN2, SOX2, ONC206, ONC212

## Abstract

Prostate cancer (PCa) is the leading cause death from cancer in men worldwide. Approximately 30% of castrate-resistant PCa’s become refractory to therapy due to neuroendocrine differentiation (NED) that is present in <1% of androgen-sensitive tumors. First-in-class imipridone ONC201/TIC10 has shown clinical activity against midline gliomas, neuroendocrine tumors and PCa. We explored the question of whether NED promotes sensitivity to imipridones ONC201 and ONC206 by inducible overexpression of SOX2 and BRN2, well-known neuroendocrine drivers, in human PCa cell lines DU145 or LNCaP. Slight protection from ONC201 or ONC206 with SOX2 and BRN2 overexpression was observed in the inducible LNCaP cells but not in the DU145 cells. At 2 months, there was an apparent increase in CLpP expression in LNCaP SOX2-overexpressing cells but this did not confer enhanced sensitivity to ONC201. DU145 SOX2-overexpressing cells had a significantly reduced ONC201 sensitivity than DU145 control cells. The results support the idea that treatment of castrate-resistant prostate cancer by imipridones may not be significantly impacted by neuroendocrine differentiation as a therapy-resistance mechanism. The results support further testing of imipridones across subtypes of androgen-sensitive and castrate-resistant prostate cancer.

## Introduction

Prostate cancer (PCa) is the leading cause of cancer-related death in men worldwide [1]. Treatment with androgen pathway inhibitors (APIs) initially suppresses cancer growth, but invariably induces castration resistance when administered to patients with metastatic disease [2]. Castration-resistant prostate cancer (CRPC) employs several documented mechanisms to evade API-induced cell death, including AR overexpression, AR mutations, and AR splice-site mutations [3]. In its most advanced and aggressive form, CRPC may adopt a neuroendocrine (NE) phenotype via the histological transformation known as neuroendocrine differentiation (NED). Patients with neuroendocrine prostate cancer (NEPC) readily develop metastases and have poor survival outcomes [4]. The standard therapy for NEPC is cytotoxic platinum chemotherapy, and there is a dire clinical need for more effective treatments [4].

The imipridone class of drugs induces cancer-specific cell death by agonizing the mitochondrial serine protease CLpP, which ultimately activates the integrated stress response via ATF4 to induce apoptosis [5, 6]. The imipridone dordaviprone (ONC201/TIC10) has shown initial efficacy in treating neuroendocrine tumors in a Phase II clinical trial [7]. The current suite of imipridones includes ONC201, ONC206, and ONC212, which feature modifications of the same pharmacological backbone and have variable potencies [6].

Previous research has elucidated NED as a transcriptional and epigenetic process, aided by genetic events such as the functional loss of p53 and Rb [8, 9]. SOX2 and BRN2 are two transcription factors implicated in driving NED, and both are commonly overexpressed in NEPC [10]. SOX2 is a pluripotency and neural transcription factor thought to activate neuroendocrine genes and suppress adenocarcinoma genes, thereby allowing redifferentiation to a new histological lineage [10, 11]. BRN2 is a neural transcription factor which activates some of the neural lineage transcriptional networks characteristic of NEPC [12, 13].

The present studies seek to evaluate the impact of BRN2 and SOX2 overexpression on two prostate cancer cell lines of different API-sensitivity status, especially with respect to imipridone sensitivity. This work serves as an initial survey of imipridone efficacy against PCa harboring aberrant expression of these transcription factors.

## Methods

### Cell Culture

LNCaP (ATCC CRL-1740), DU145 (ATCC HTB-81), PC3 (ATCC CRL 1435), and 22Rv1 (ATTC CRL-2505) prostate cancer cell lines were cultured in RPMI-1640 (Cytiva SH30027.FS) with 10% fetal bovine serum (FBS) and 1% penicillin-streptomycin (PS). Inducible cell lines were cultured in the same media, but with tetracycline-free FBS. NEPC cell line NCI-H660 (ATTC CRL-5813) was cultured in DMEM (Cytiva SH30022.02) +20% FBS and 1% PS. NK92-MI cells were cultured in alpha-MEM +10% FBS, 10% horse serum, 1% PS, 1% non-essential amino acids, 1% L-glutamine, 1% Sodium Pyruvate, 1% myoinositol, 0.1mM β-mercaptoethanol, and 2mM folic acid. All cells were incubated at 37ºC and 5% CO2.

### Inducible Cell Line Generation

Vector and insert DNA was linearized from the pLX-TRE-dCas9-KRAB-MeCP2-BSD lentiviral plasmid (Addgene 140690) and SOX2 or POU3F2 pcDNA3.1(+)-P2A plasmids ordered from Genscript. Linearization was performed by PCR with PrimeStar Max DNA polymerase (Takara R045A) per the manufacturer’s protocol. Vector and insert DNA were then purified by agarose gel electrophoresis followed by gel extraction. Lentiviral plasmids were generated from the linearized fragments using the Takara In-Fusion Cloning kit (Takara 638947) to create pLX-TRE-EV-BSD, pLX-TRE-SOX2-BSD, and pLX-TRE-BRN2-BSD lentiviral plasmids. All lentiviral plasmids were transformed into NEBStable competent E coli.

For lentivirus preparation, five million HEK293T cells (ATCC CRL-3216) were seeded in a 10 cm tissue culture dish in antibiotic-free DMEM with 10% FBS (ATCC 30-2020). After adhering overnight, they were transfected with 50 µL of Lipofectamine 2000 (ThermoFisher #11668019); 10 µg of pLX-TRE-SOX2-BSD, pLX-TRE-BRN2-BSD, or pLX-TRE-EV-BSD transfer plasmid; 5 µg of pMDLg/pRRE (Addgene 12251, pMDLg/pRRE was a gift from Didier Trono); 5 µg pRSV-Rev (Addgene 12253, pRSV-Rev was a gift from Didier Trono); and 2.5 µg of pMD2.G (Addgene 12259, pMD2.G was a gift from Didier Trono). The medium was replaced after 16 hours. 48 hours after transfection, the medium was harvested, centrifuged at 500g for 5 minutes, and the supernatant was filtered through a 0.45 µm polyethersulfone syringe filter (Millipore SLHPR33RS).

LNCaP and DU145 were plated at 50% confluency in a 12-well plate and incubated overnight. The cells were then exposed to lentivirus for three days and subsequently selected with blasticidin (4.5 µg/mL LNCaP, 4 µg/mL Du145) for 14 days. After selection, inducible cells were consistently cultured in 2 μ g/mL blasticidin. For experiments, induction was performed with 100ng/mL doxycycline (DOX) in medium without blasticidin.

### Western Blots

A total of 400,000 cells were plated in 12 or 6 well tissue culture plates and allowed to adhere overnight before treatment. After treatment or overexpression, cells were harvested and immediately lysed with RIPA buffer (Sigma-Aldrich R0278) including a protease inhibitor cocktail (Sigma-Aldrich 04693159001). SDS was added to denature the proteins, which were boiled at 95ºC for 10 minutes. Protein amounts (5-20 µg depending on the gel) were normalized by BCA assay and run on SDS-PAGE gels (Invitrogen), before being wet-transferred to a PVDF membrane. The membrane was blocked for 1 hour in 5% wt/vol milk in TBST before being incubated overnight in primary antibody. Primary antibodies used include vinculin (Cell Signaling 4650), β-actin (Cell Signaling 4970), SOX2 (Cell Signaling 2748S), BRN2 (Cell Signaling 12137S), CgA (Abcam ab283265), DR5 (Cell Signaling 3696S), CLpP (Cell Signaling 14181S), PGP9.5 (Cell Signaling 3524), syp (Cell Signaling 5461), H3K27me3 (Cell Signaling 9733), and Cyclin D1 (Cell Signaling 2978S). The next day, the membranes were washed three times for 10 minutes in TBST, followed by 1-hour incubation in the corresponding HRP-conjugated anti-mouse (Thermo Scientific 31430) or anti-rabbit (Thermo Scientific 31460) secondary antibody. The membranes were again washed with TBST three times for 10 minutes. The membranes were subsequently imaged using Pierce™ ECL (ThermoFisher 32106), SuperSignal™ West Pico PLUS (ThermoFisher 34580), or SuperSignal™ West Femto (ThermoFisher 34096) chemiluminescent substrates with the Syngene imaging system and software.

### Cell Viability Assays

LNCaP and DU145 were plated at 5,000 cells per well and NCI-H660 was plated at 8,000 cells per well in 100 µM media in a 96-well plate. Cells were allowed 24h to adhere before being treated in triplicate. The CellTiter-Glo® assay (Promega G9241) was used to assess viability.

### Scratch Assays

Inducible cells were plated in triplicate at 60,000 cells per well in a 96-well plate with DOX and allowed to adhere overnight. The following day, cells were scratched with a sterile 2 µL plastic pipette tip and the wells were washed with PBS. Cells were imaged daily for 8 days using the brightfield channel of the ImageXpress HT.ai confocal microscope. Images were montaged and analyzed with the Wound Healing Assay plugin for Fiji [14]. Statistical significance of width and area measurements over time and overexpression condition were determined via two-way ANOVA, followed by unpaired t-test with Tukey’s correction for multiple comparisons.

### Clonogenic and Cell Growth Assays

DU145 inducible cells were plated at 300 cells per well in a 12-well plate with DOX in triplicate. The following day, 1 µM ONC201, 3 µM ONC201, or DMSO was added to the wells. The cells were treated for 8 days before fixing and staining. LNCaP inducible cells were used in a higher density cell growth assay, and were plated at 10,000 cells per well in a 12-well plate with DOX in triplicate. The following day, ONC201 was added in doses of 0.75µM, 1µM, or 4µM and ONC206 was added in concentrations of 100nM, 200nM, or 500nM. A volume of DMSO corresponding to the highest dose of ONC201 or ONC206 was used to control for the respective drug. LNCaP inducible cells were treated for 5 days. After treatment, the wells were washed with PBS, fixed with methanol, and dyed with Coomassie blue. After washing, plates were imaged with the Syngene imaging system and analyzed with the ColonyArea plugin for Fiji with automatic thresholding [15]. Statistical significance was determined via two-way ANOVA, followed by unpaired t-test with Tukey’s correction for multiple comparisons.

## Results

### Characterization of an inducible model for SOX2 and BRN2 overexpression in LNCaP and DU145

After 24h of DOX induction, western blots from LNCaP and DU145 cells infected with pLX-TRE-EV-BSD (EV), pLX-TRE-SOX2-BSD (iSOX2), or pLX-TRE-BRN2-BSD (iBRN2) revealed protein-level upregulation of the intended transcription factor (**Figure 1A**). Western blots after 4d of DOX showed a robust increase in the NE marker synaptophysin (syp) in iBRN2 cell lines (**Figure 1B**) There appeared to be marginal increases in the NE marker chromogranin A (CgA) protein in LNCaP iBRN2 and DU145 iSOX2. The NE marker NCAM1 was not detectable by western blot. iBRN2 or iSOX2 did not detectably alter the protein expression of one another in any cell line tested at 4d. There were no clear changes in the expression of ONC201 target CLpP or its chaperone CLPX.

**Figure 1.**
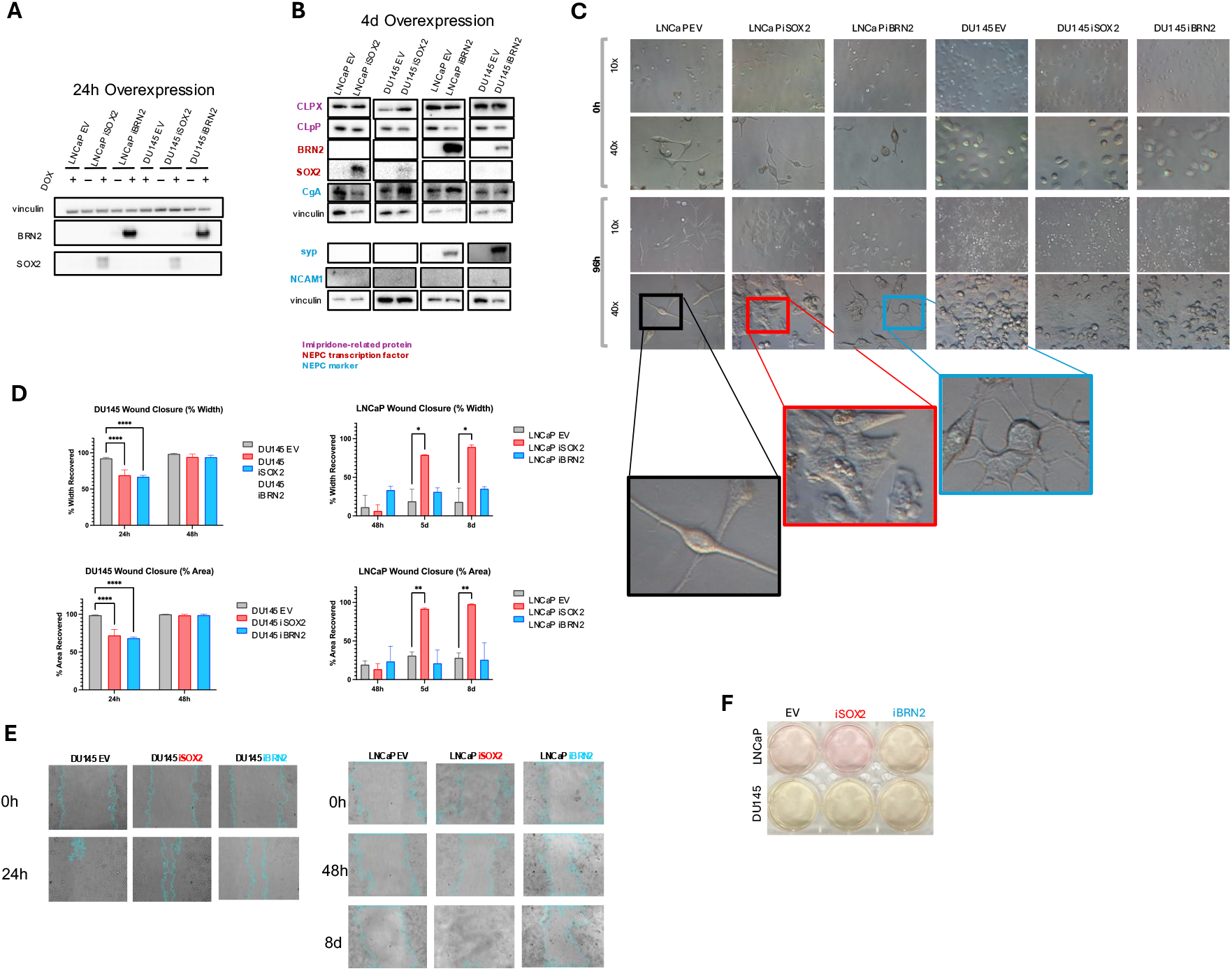
Characterization of NE-driver inducible system in LNCaP and DU145. **(A)** Western blot for the overexpressed transcription factors in all inducible cells engineered. DOX-negative conditions are included to evaluate leak in the system. **(B)** Western blot of the overexpressed transcription factors (red), neuroendocrine markers (blue), and ONC-related proteins (purple) after 4 days of DOX induction. **(C)** Images of inducible cells after at 0 and 4 days of induction. Objective is noted to the left of the images. **(D)** Normalized quantification of scratch assays for DU145 and LNCaP inducible cell lines after 4 days of DOX induction without treatment. Percent width and area values are reported at 24 and 48 hours for DU145 and 48 hours, 5 days, and 8 days for LNCaP. **(E)** Representative images of scratch assays with FIJI-generated migration borders indicated by thin cyan lines. **(F)** Media color for inducible cell lines after 4 days of DOX induction.

Images after 4 days of induction reveal morphological changes in LNCaP cells, with both iSOX2 and iBRN2 cells losing the stellate shape characteristic of LNCaP (**Figure 1C**). Changes in pH were noted by differences in media color after 4 days depending on the cell line, with LNCaP iSOX2 being less acidic and LNCaP iBRN2 being more acidic than the EV (**Figure 1F**). This trend was not present for the DU145 cell lines.

The migratory potential of inducible cell lines was determined by scratch assay. Interestingly, SOX2 and BRN2 overexpression hindered wound healing in DU145 (**Figures 1D, E**). A reverse effect was observed with iSOX2 in LNCaP, as iSOX2 greatly enhanced wound-healing. LNCaP is not an ideal cell line for a scratch assay due to its weak adherence [16]. It was also noted that LNCaP iSOX2 adhered less tightly to plates than LNCaP EV or LNCaP iBRN2, posing a potential confounding variable for the observed increased migratory potential LNCaP iSOX2.

### Short-term overexpression of SOX2 and BRN2 alters imipridone efficacy

To assess the sensitivity of these cell lines to imipridones, they were induced with DOX for 4 days, adhered for 24h, and treated with ONC201 or ONC206 for 72h in DOX. LNCaP iSOX2 appeared to have the greatest change in sensitivity to both imipridones tested, as the viability plateaued at doses above the IC50 drug concentration without approaching 0% (**Figure 2A**). This effect was also present with LNCaP iBRN2, but to a lesser extent. In the inducible DU145 cell lines, similar trends were observed in imipridone sensitivity, but not to the same magnitude. Changes in IC50 were minor with overexpression, and thus, area under the curve (AUC) was calculated to compare sensitivity quantitatively (**Figure 2A**) Both iSOX2 cell lines had significantly larger AUC than the

**Figure 2.**
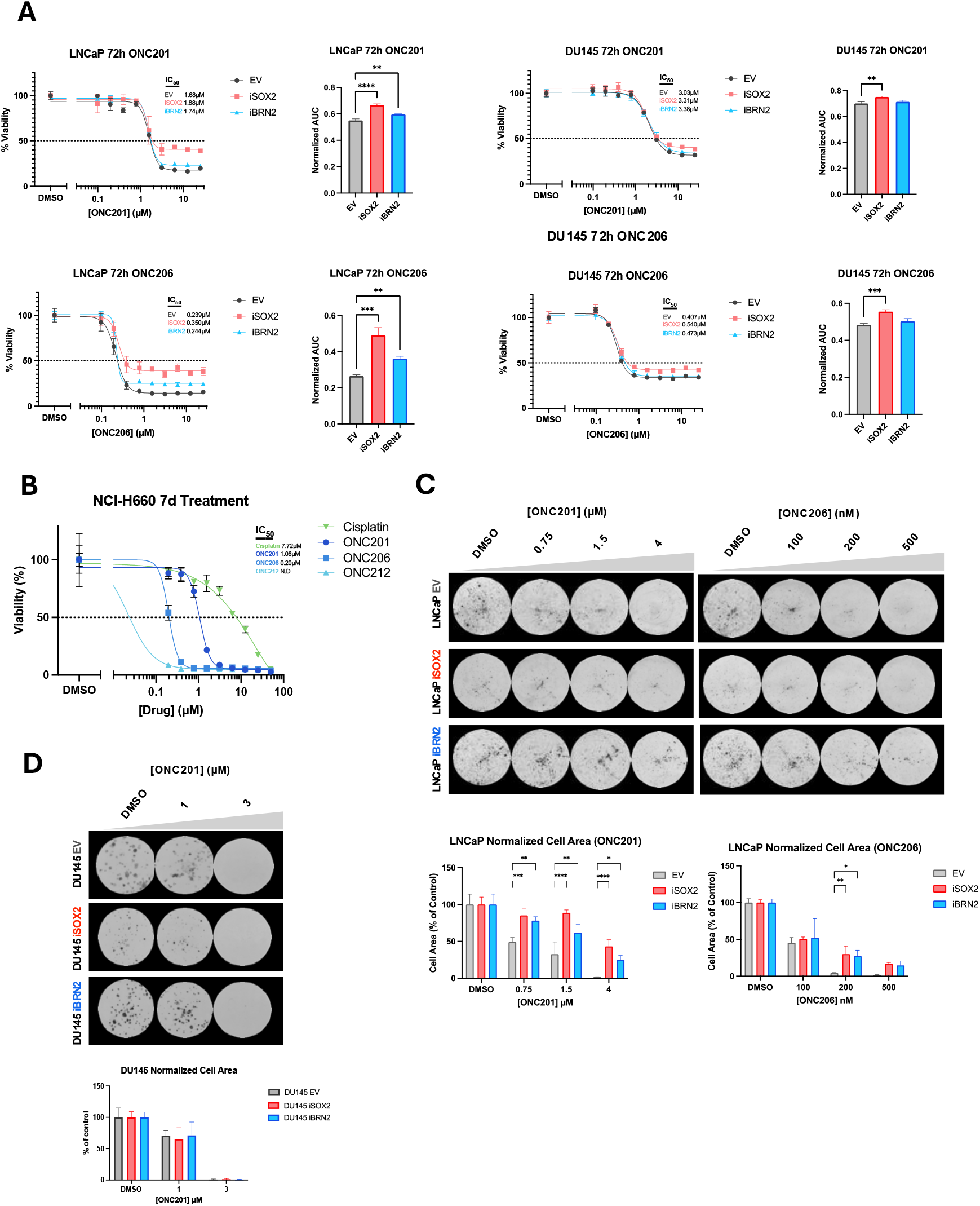
Characterization of NE-driver inducible system in response to imipridones. **(A)** Dose-response curves for LNCaP EV, iSOX2, and iBRN2 and DU145 EV, iSOX2, and iBRN2 treated with ONC201 or ONC206 for 72h after 4d DOX induction. Area under the curve for dose-response curves are to the right of each curve, with 1 defined as the area from the lowest imipridone dose to the highest, excluding the DMSO condition. **(B)** Dose-response curves for NE cell line NCI-H660 treated for 7d with ONC201, ONC206, ONC212, or cisplatin for 72h. **(C)** Representative images and area quantification of LNCaP cell growth assay in ONC201 or ONC206 after 4 days DOX induction. Cells were fixed, stained, and imaged after 5 days of treatment. **(D)** Representative images and area quantification of DU145 colony formation assay in ONC201 after 4 days DOX induction. Cells were fixed, stained, and imaged after 8 days of treatment.

EVs, indicating decreased drug sensitivity. BRN2 induction only significantly increased the AUC for LNCaP, though the mean AUC for DU145 iBRN2 was still higher than the EV. To test the efficacy of imipridones on a bona fide NEPC cell line, NCI-H660 was treated for 7d with ONC201, ONC206, or ONC212, all of which were more potent than cisplatin (**Figure 2B**). Due to its low capacity to form colonies, LNCaP cells were used in a higher-density growth assay, where they were treated with ONC201 or ONC206 for 5d following 4d induction (**Figure 2C)**. LNCaP iSOX2 and iBRN2 generally exhibited higher resistance to ONC201 and ONC206 relative to the EV. While the number of cells plated between overexpression conditions remained constant, LNCaP iSOX2 appeared to have a lower baseline growth potential, with less area than iBRN2 or EV in the untreated condition. The LNCaP iSOX2 cells that grew were relatively resistant to the imipridones, as evidenced by small decreases in area with escalating drug doses.

In DU145, clonogenic assays were used to determine if the differential responses to ONC201 persisted over time, but there were no significant differences in the cell area or intensity after 8 days (**Figure 2D**).

### Long-term overexpression of SOX2 and BRN2 does not mimic the changes observed with short-term overexpression

LNCaP and DU145 cells were cultured in DOX for 2 months before being assessed for their sensitivity to ONC201. Overexpressed TFs were still detectible by western blot (**Figure 3A**). Notably, there was an apparent increase in CLpP expression in LNCaP iSOX2. This increase did not confer enhanced sensitivity to ONC201, as there were no significant changes in AUC (**Figure 3B, C**). DU145 iSOX2 had a significantly higher AUC than DU145 EV.

**Figure 3.**
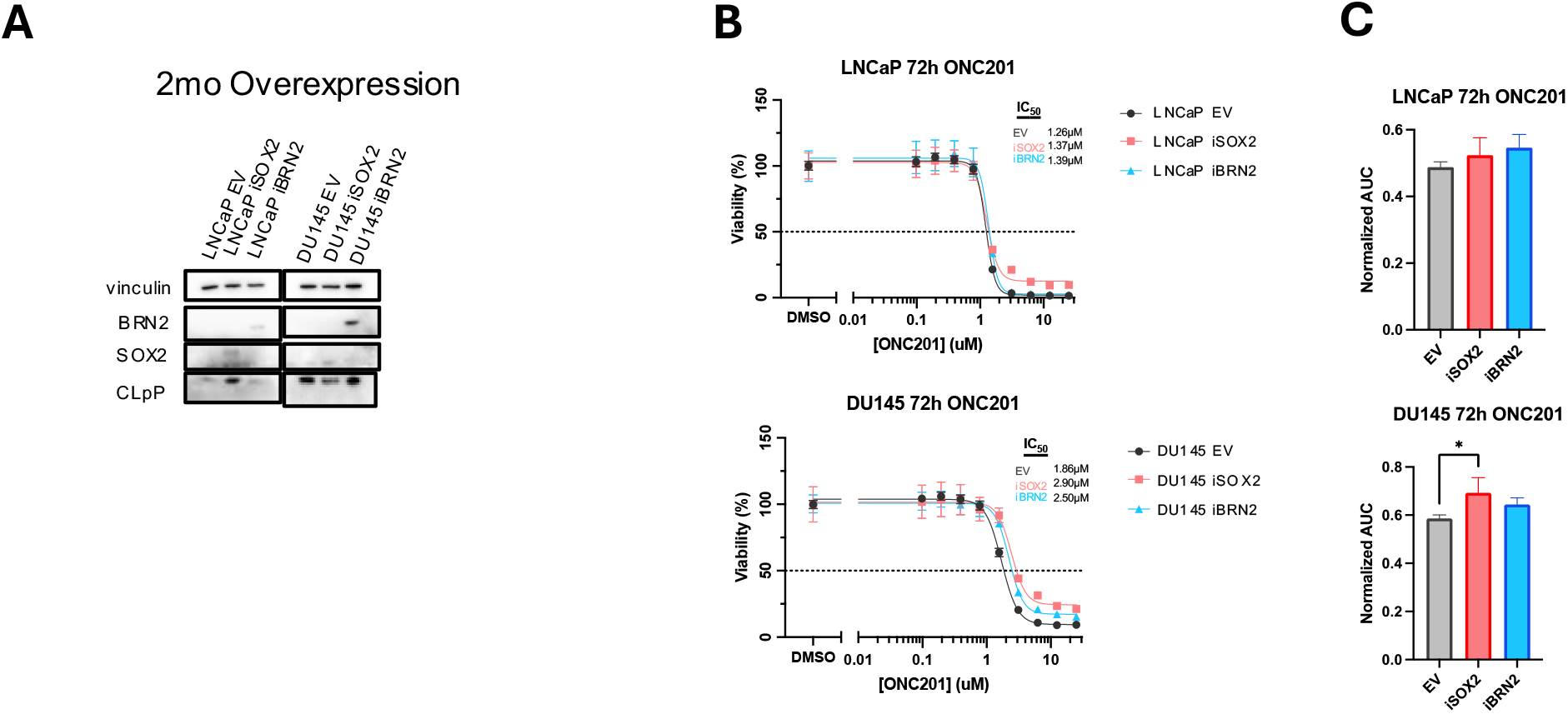
Sustained overexpression of BRN2 and SOX2 alters response to ONC201. **(A)** Western blot for the overexpressed transcription factors and CLpP in all inducible cells engineered. **(B)** Dose-response curves for 72h ONC201 treatment after two months of DOX induction. **(C)** Area under the curve for dose-response curves, with 1 defined as the area from the lowest imipridone dose to the highest, excluding the DMSO condition.

## Discussion

Neuroendocrine differentiation (NED) continues to be an elusive resistance mechanism to API therapy. Identifying and exploiting vulnerabilities in NEPC or the NED process presents an opportunity to improve clinical outcomes. As an emerging class of therapeutics, imipridones have proven efficacious in targeting various tumor types, warranting an investigation of their efficacy against NEPC. Here, we examine two purported drivers of NED in SOX2 and BRN2 and evaluate the effects of their induction in the LNCaP and DU145 cell lines concerning NE marker expression, migration, and imipridone sensitivity. LNCaP and DU145 were selected for the inducible NE-driver model for their distinct genetic and phenotypic characteristics. LNCaP is a standard castration-sensitive prostate adenocarcinoma model for in vitro studies and xenografts, as it expresses AR, PSA, and wild-type p53 and Rb [17, 18]. An inducible model for SOX2 overexpression in LNCaP has been described previously by Metz et al in 2020 [19], but to our knowledge, this is the first inducible BRN2 model in the cell line. In contrast to LNCaP, DU145 is commonly referred to as CRPC–though not NEPC–due to its lack of AR expression, and mutations in TP53 and RB1 [17, 20]. Thus, overexpressing SOX2 and BRN2 in DU145 was an attempt to push the cell line further down the spectrum toward NEPC. To our knowledge, such models have not been previously engineered in DU145.

We recreated some aspects of the NE phenotype through transcriptional reprogramming, such as synaptophysin expression with iBRN2. This effect was apparent in both LNCaP and DU145, expanding upon previous findings in the literature. A seminal paper on the role of BRN2 in NEPC used LNCaP mouse xenografts treated with the API enzalutamide to develop resistant cells [13]. In this work, the authors found BRN2 to be essential for NEPC marker expression, reported direct repression of BRN2 by AR, and determined that exclusively overexpressing BRN2 *or* SOX2 would increase the mRNA expression of one another. While our results did corroborate these mutual changes in SOX2 and BRN2 expression at the protein level or the reported increases in CgA and NCAM1, the observed increase in synaptophysin was consistent in both studies. That said, it is important to consider that the prior work used a different model from the present study, in which overexpression was performed in API-naïve cells. Contrasting with the observations of Metz et al [19], iSOX2 did not detectibly increase synaptophysin in our LNCaP model. Differing from Metz et al, our LNCaP EV cell line did not express a base level of synaptophysin, suggesting differences between the two models that may be the basis of the observed differences.

Experiments with imipridones indicate that SOX2 and BRN2 do not make prostate cancer cell lines more sensitive to imipridone treatment. On the contrary, initial differences in imipridone sensitivity at 72h depending on SOX2 and BRN2 overexpression indicated more resistance in iSOX2 and iBRN2 cell lines. These differences were not apparent at 8d using a clonogenic assay in DU145. A possible explanation for the initial observed differences could be altered growth rates between the cell lines. For instance, an extended doubling time in iSOX2 cell lines could have made these cells appear more resistant to treatment. The apparent increase in CLpP in LNCaP iSOX2 after 2 months DOX agrees with past research describing increases in mitochondrial count in prostate cancer cells with heightened SOX2 expression [21]. That said, the present CLpP increase was not accompanied by imipridone sensitivity, suggesting the existence of other molecular modulations in the cell offsetting the increase in the imipridone target.

Recent research has uncovered heterogeneity in the NED process, with subpopulations of cells expressing differential transcription factor profiles after reprogramming [22, 23]. The myriad of transcriptional drivers and markers associated with NEPC imply that development of the disease may not be limited to a single transcriptional pathway. Further, the clinical development of NEPC is complicated by a number of factors including tumor microenvironment, hormone levels, and immune interactions [24, 25]. These in-vitro experiments do not recreate the environment in which NED develops in patients. For one, the experiments were not carried out with functional inhibition of AR using APIs or androgen-free media. Additionally, the LNCaP cells where the highest degree of change in imipridone response was observed have wild-type p53 and Rb, two tumor suppressor proteins that are almost always altered in the development of NED. Further research must gain insight to the roles of these tumor suppressors in NED.

Continuing to build an understanding of the molecular events on the path to NED will allow for the development of more efficacious therapies. Simultaneously, in the dawning era of personalized medicine, advances in sequencing technologies will help identify cases in which specific emerging therapeutics will provide the greatest clinical benefit, leading to improved patient outcomes. While short-term SOX2 and BRN2 induction provided some protection from imipridones, it did not confer complete resistance, as cells were still killed by marginally higher doses. The present study found no increase in sensitivity, given the heterogeneous and complex nature of the disease, it remains to be determined whether imipridones will play a role in the future of NEPC treatment.

## Acknowledgments

W.S.E-D. is an American Cancer Society Research Professor and is supported by the Mencoff Family University Professorship at Brown University. This work was supported by an NIH grant (CA173453) to W.S.E-D. The contents of this manuscript are solely the responsibility of the authors and do not necessarily represent the official views of the National Cancer Institute, the National Institutes of Health, or the American Cancer Society.

